# Spatial variations in sap flow rates in mature tree stems before and after drilling treatment

**DOI:** 10.1101/2024.10.07.616919

**Authors:** Toshihiro Umebayashi, Taku M. Saitoh, Kenji Tsuruta, Dennis Otieno

## Abstract

The dye uptake and heat transfer methods are used to understand sap movement in mature tree stems, but correlation of results between both methods is weak. The circumferential variation of sap movement is detected generally in the heated transfer method. On the other hand, the tangential movement such as helical ascent is tracked in dye uptake method. It is still unclear whether the circumferential and the tangential movements in both methods mean same and/or related phenomenon. In this study, we monitored sap flux density (*F*_d_) on the east and west sides in intact stems of *Betula ermanii* Cham. using the thermal-dissipation method to identify *F*_d_ response to the creation of artificial flows through severed vessels by drill treatments. We then tracked sap flow by injecting dye solution. The *F*_d_ values in east side tended to be higher than that in west side before drilling treatment. Similar *F*_d_ values in both sides were detected after drilling treatments, and dye heights were similar to *F*_d_ values after drilling in any sides. Simple comparison of water ascent using both methods can lead to errors, because water ascent from severed vessels may differ from water ascent via roots.

## Introduction

Water in terrestrial plants generally moves from the roots to the leaves under negative pressure created by transpiration. Recent studies have investigated the water distribution in intact xylem conduits (tracheids or vessels) at the cellular level using micro-computed tomography (micro-CT; Brodersen et al. 2010; Choat et al. 2016), magnetic resonance imaging (MRI; Holbrook et al. 2001), and cryo-scanning electron microscopy (Utsumi et al. 1996; Umebayashi et al. 2016), and have visualized the three-dimensional structure of xylem conduits at the cellular level using confocal laser scanning microscopy (Kitin et al. 2004) and micro-CT (Brodersen et al. 2013; Choat et al. 2015; Torres-Ruiz et al. 2015). But non-invasive imaging of water movement has rarely been reported. Most of existing literature also refer to saplings and cut branches.

Thermal imaging of the ascent of sap (Anfodillo et al. 1993) and flow MRI (Windt et al. 2006) are useful in studying water movement in intact plants, but they are not common. Instead, the dye uptake method has been regularly used for detailed investigation of water movement in many woody plants (Baker and James 1933; Sano et al. 2005; Umebayashi et al. 2007, 2008, 2010). Vessel plumbing in stems of mature trees is complex as demonstrated by dye experiments, showing varying patterns of dye ascent such as helix or straight (Baker and James 1933; Kozlowski and Winget 1963, Umebayashi et al. 2008, 2010). This method is, however, destructive and may introduce artifacts on opening conduits underwater with a saw or a drill before the introduction of dye to stain the conduits (Wheeler et al. 2013).

Heat transfer methods which monitor the thermal loss caused by sap flow, such as the heat-pulse velocity, heat balance, thermal-dissipation (TD), and heat field-deformation methods (e.g., Čermák and Nadezhdina 1998; James et al. 2002; Gartner and Meinzer 2005) also show radial and azimuthal (i.e. circumferential) variations of water movement in intact trees. Especially, in TD method which has been widely used to estimate the water movement because of its low cost and acceptable accuracy, circumferential variations are well detected not only in conifer (Tsuruta et al. 2010), but also in broadleaved trees (Lu et al. 2000; Tateishi et al. 2008). Circumferential variation of *F*_d_ values observed with the TD method may be induced by growth conditions in the field such as partial shading of the tree crown (Oren et al. 1999) or by root distribution and water availability in the soil (Lu et al., 2000).

However, within the stem of a single tree, correspondence between *F*_d_ measured using the TD method and dye height is weak (Tsuruta et al. 2010). Čermák et al. (1992) measured the xylem sap flow rate by the heat balance method, and explained that the introduction of severed conduits causes changes in xylem pressure, which dramatically increase *F*_d_. This result suggests that artifacts of the dye uptake method can explain by monitoring *F*_d_ before and after the introduction of severed conduits. The variance between the two methods, however, has been considered when detecting water ascent in sapwood using dye uptake method (Tateishi et al. 2008; Tsuruta et al. 2010). Thus, we need to understand the cause of the variance between *F*_d_ measured using the TD method and dye height by combining both techniques.

To explain the characteristic of water flow from severed conduits and to reveal artifacts of the dye uptake method, especially how the opening of conduits eliminates circumferential variation, we examined water movement in intact main stem of a common birch species with diffuse-porous wood (*Betula ermanii* Cham.). We monitored the change of *F*_d_ before and after the creations of severed conduits using TD method. We drilled four holes below each TD sensor under water to induce changes in *F*_d_. Then we used the dye-uptake method to track sap flow from the drill holes, and compared dye ascent and *F*_d_ on the east and west sides of the stems.

## Materials and methods

Sap flux density (*F*_d_, g m^−2^ s^−1^) was monitored in two *B. ermanii* trees growing at Gifu University, Takayama (36°08′N, 137°25′E, 1420 m a.s.l.), using the TD method. The mean annual air temperature and the total precipitation at the nearest field station (500 m south of Takayama) were 6.5 °C and 2089 mm (average from 1996 to 2009). The two trees (b1 and b2) were 18.0 and 17.1 m tall, and their diameters at breast height (DBHs) were 17.3 and 16.7 cm, respectively. *F*_d_ was measured on the east and west sides of the stems on 3 and 4 August 2011.

The TD sensor consisted of two thermocouple probes 20 mm long and 2 mm wide. Each probe included a 0.2-mm-diameter copper–constantan thermocouple. The two thermocouples were joined at the constantan leads so that the voltage measured across the copper leads provided the temperature difference between the upper (heat) probe and the lower (reference) probe. Both probes were inserted, and the upper probe was set about 15 cm above the lower probe (Fig. S1). Both were covered with a radiation shield to minimize the direct thermal load.

To understand the spatial distribution of water movement in intact stems, we monitored *F*_d_ before and after the creation of severed vessels below each TD sensor by drilling the stem under water. The sensors pairs were installed at depths of 0 to 20 mm in the sapwood to monitor *F*_d_ of the outer xylem. Each upper probe was set at 1.3 m above the ground and received a constant 0.2 W to heat the sapwood. The temperature difference between the two probes (Δ*T*) was scanned every 5 s, with values recorded as 1-min means (CR1000, Campbell Scientific, Logan, UT, USA) to detect any changes in *F*_d_ before and after water movement from the severed vessels that were made under water. *F*_d_ was calculated from Δ*T* by the method of Granier (1987) on the assumption of zero *F*_d_ (i.e., Δ*T*_max_) at night as *F*_d_ = 119 [(Δ*T*_max_ – Δ*T*) / Δ*T*]^1.231^.

Drill treatments for the dye uptake method were started at 12:10 on 4 August 2011 (Table 1). On each side of a tree (west and east), a 6-cm horizontal line was drawn 20 cm below the reference sensor probe (Fig. S1). A plastic bowl was attached to the stem below this line with caulking material and was filled with distilled water. Holes were then carefully drilled along the line, under the water. Four holes (each 9 mm in diameter, about 15 mm apart) were drilled from left to right along each line. Holes were drilled to the half of the stem diameter, and there was a delay of at least 10 s between drilling of consecutive holes to confirm that the *F*_d_ changed on the display of a PC connected to the datalogger. Holes were drilled also on the north side of each tree to explore the dye height.

**Table 1.** Start times of underwater drilling and dye uptake, and maximum dye height in *Betula ermanii* trees b1 and b2.

| Sample No. | Height (m) | DBH (cm) | Stained sapwood thickness (cm) | Side of stem drilled | Start time of drilling | Start time of dye uptake | Max. dye height (m) |
| --- | --- | --- | --- | --- | --- | --- | --- |
| b1 | 17.1 | 17.3 | 3.5 ± 0.3<sup>a</sup> | east | 12:10 | 12:50 | 0.50 |
|  |  |  |  | west | 12:18 |  | 0.50 |
|  |  |  |  | north |  |  | 0.50 |
| b2 | 18.0 | 16.7 | 4.0 ± 0.3<sup>a</sup> | east | 12:37 | 13:15 | 0.75 |
|  |  |  |  | west | 12:45 |  | 0.50 |
|  |  |  |  | north |  |  | 0.75 |

<sup>a</sup> n=3, mean ± standard deviation

After the holes were drilled, the water in the bowl was replaced with a dye solution (0.5% acid fuchsin filtered to 1 µm). After perfusion for 30 min, the tree was cut down below the drilling point. Discs 3 cm thick were collected at 0, 10, 30, and 50 cm above the dye uptake point, and at 25-cm intervals above 50 cm to the maximum height of the stained xylem.

Segments with stain were immediately stored in liquid nitrogen. In addition, the dye distribution at the sites of all sensors was confirmed and photographed along the transverse surface. All samples were transported in dry ice to the University of Tokyo and then freeze-dried at −50 °C for 48 h (FDU-810 freeze dryer, Eyela, Tokyo, Japan) for analysis. Stained areas were identified under a stereomicroscope.

We measured several meteorological components at the nearest field station (500 m south of Takayama site). Short-wave downward radiation was measured with solar radiation radiometers (CMP3, Kipp & Zonen, Delft, the Netherlands) installed at the height of 2.8 m above ground level. Air temperature and humidity were measured at the height of 2.7 m above ground level using humidity and temperature probes (HMP45A, Vaisala, Vantaa, Finland) with ventilated radiation shields. VPD was calculated by both air temperature and relative humidity data. All these data were collected at 10-min intervals using a CR1000 datalogger (Campbell Scientific, Logan, UT).

Another two trees (DBH ≈ 5 cm) were used to visualize the water distribution around a probe according to Umebayashi et al. (2016). At midday on 3 August 2011, we drilled a sensor hole (20 mm long, 2 mm wide) in air at a height of 100 cm. A plastic collar attached before sunrise was filled with liquid nitrogen for 5 min to freeze the stem. Frozen samples were immediately excised with a handsaw and transported in dry ice to the University of Tokyo. To visualize the water distribution, we sectioned the frozen blocks on a sliding microtome, and transferred the sections to the cold stage of a cryo-SEM (JSM-6390LV, JEOL, Tokyo, Japan). The frozen cut surfaces were etched and observed uncoated at 3 kV.

## Results

Figure 1 shows the diurnal courses of solar radiation (*R*_s_), vapor pressure deficit (VPD) (Fig. 1a), and *F*_d_ at 0 to 20 mm depth in both trees on 4 August (Fig. 1b, c). Values of *R*_s_ and VPD were high around 11:50 (Fig. 1a). The peak *F*_d_ was recorded on the east side of each tree at 11:45 (Fig. 1b, c). In both trees, *F*_d_ was near 0 at midnight, and no drastic changes were observed during the nighttime. In both trees, *F*_d_ values in east side tended to be higher than that in west side before drilling, and *F*_d_ showed a steep peak after drilling (Fig. 1b, c arrows). The peak showed generally after the second or third hole was drilled, and the sudden increase in *F*_d_ stopped within 10 s after a hole was drilled. *F*_d_ then decreased dramatically. Although *F*_d_ values in both trees differed between sides before drilling, they were similar after drilling (Fig. 1b, c).

**Figure 1.**
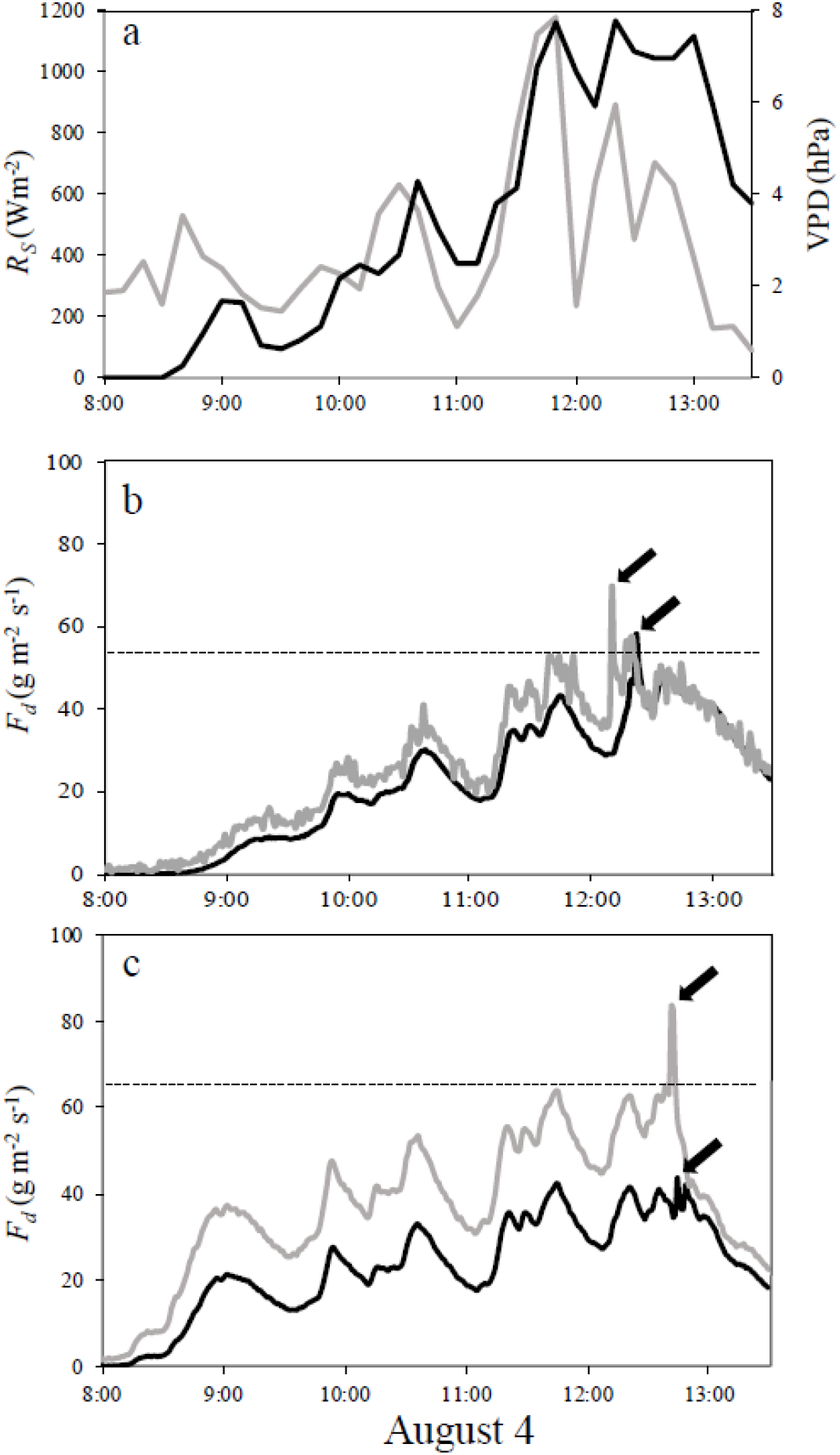
Diurnal patterns of (a) solar radiation (*R*_s_), and vapor-pressure deficit (VPD), and (b, c) sap flux density (*F*_d_) in *Betula ermanii* on 4 August 2011 in trees (b) b1 and (c) b2. (a) Black, VPD; gray, *R*_s_. (b, c) Sides of the tree: gray = east, black = west. The arrowheads and horizontal dashed lines mark the times and value of the peak before drill treatments on the west side of b1 and on the east side of b2. Arrows show the peak *F*_d_ after drilling.

The maximum height of the dye ascent was 50 cm (1.0 m h^–1^) in b1 and 50 or 75 cm (1.0 or 1.5 m h^–1^) in b2 (Table 1). In addition, the dye heights related to *F*_d_ not before, but after drilling (Fig. 2). In the tangential movement of stain within the annual rings (i.e., helical movement), the stained areas from each drill hole were distinct near the point of drilling holes (Fig. 3a, b arrows), but they merged as the dye rose (Fig. 3c). Dye was clearly observed adjacent to the probe hole (Fig. 3d).

**Figure 2.**
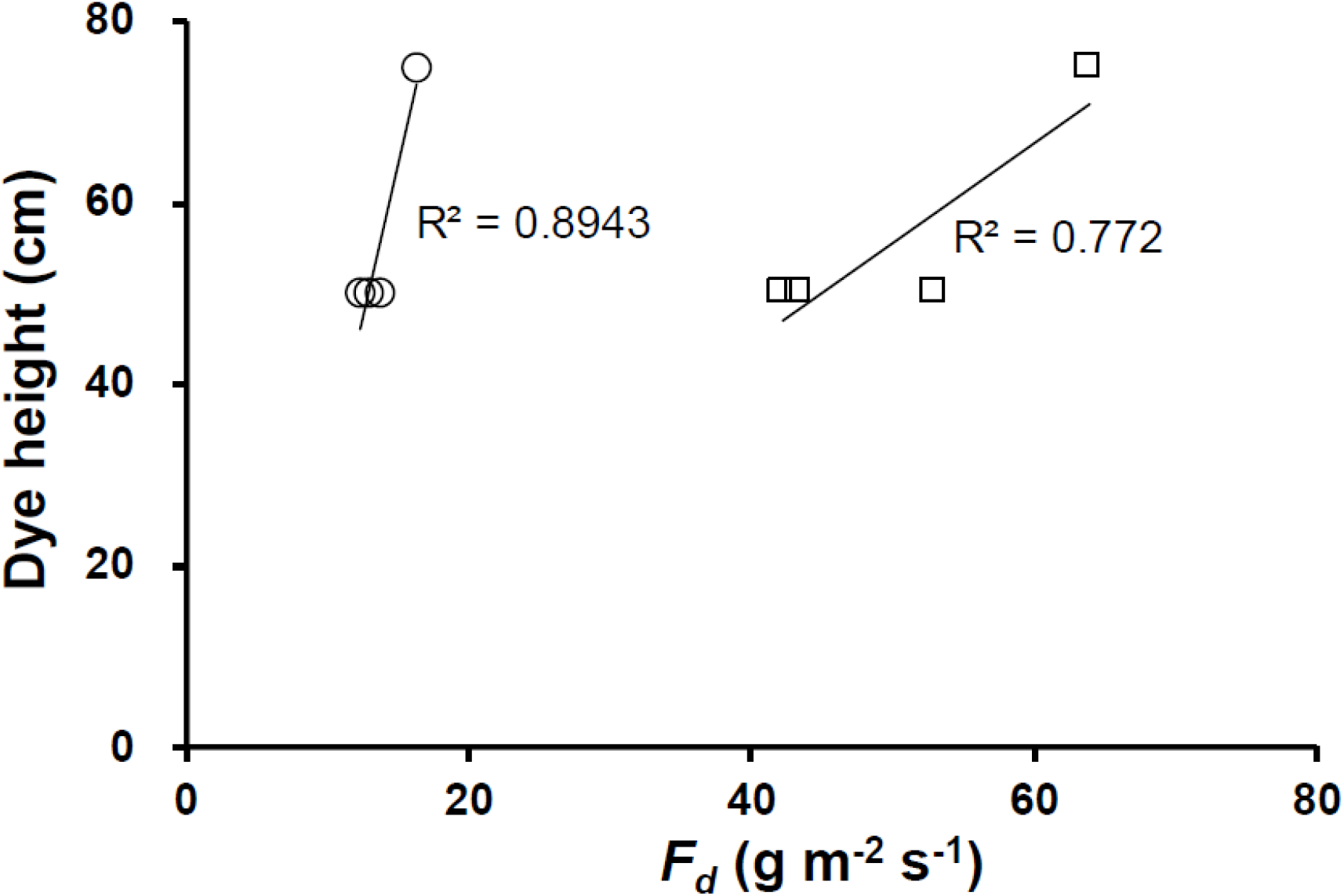
The relationship between the maximal dye height and the sap flow density (*F*_*d*_) before and after drilling treatments in each site. Square symbol shows *F*_*d*_ at 11:45, and circle symbol shows *F*_*d*_ at 13:35. Lines show each liner regression.

**Figure 3.**
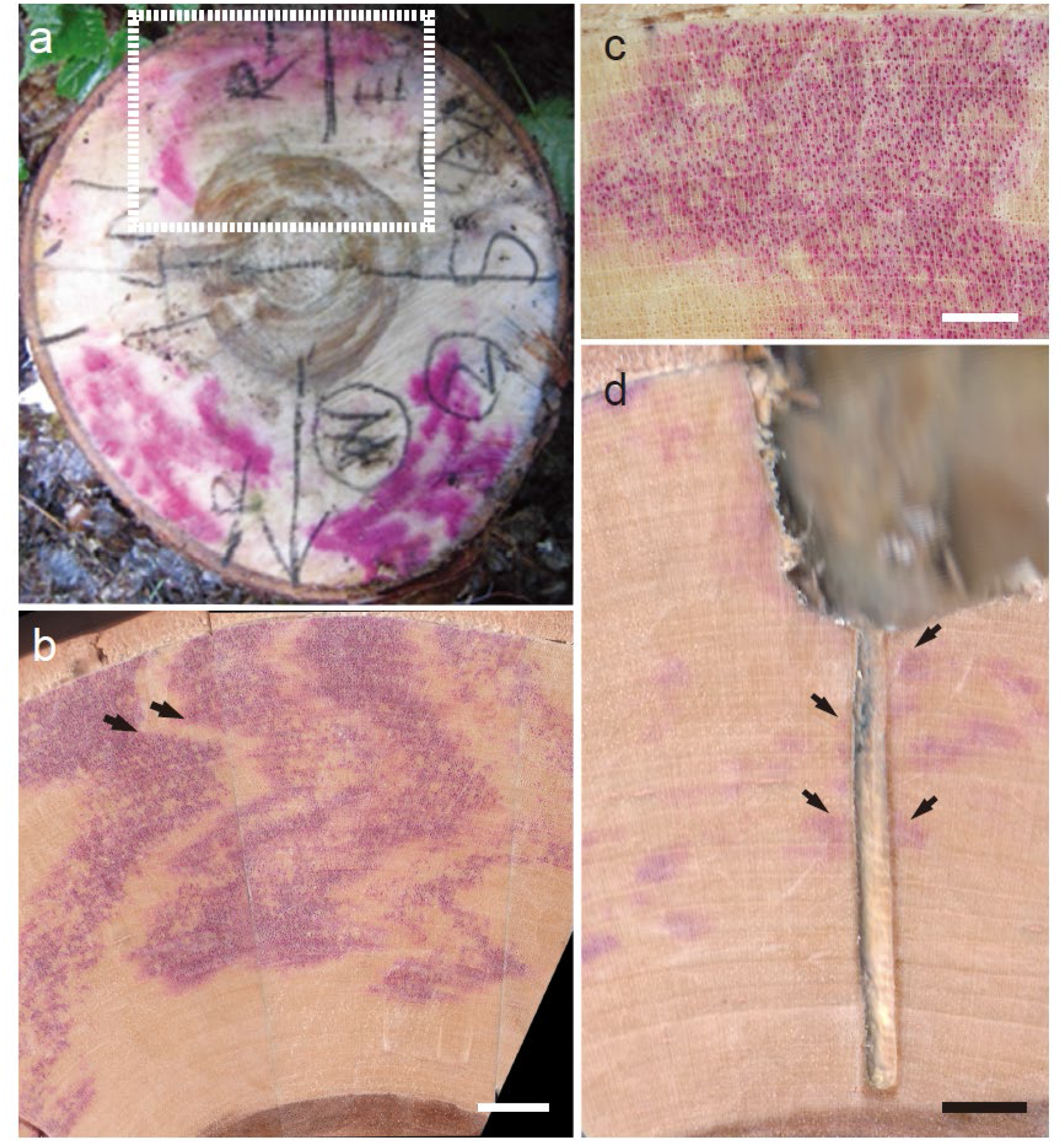
Transverse surfaces of stained xylem in *Betula ermanii* tree b2. (a) A disc near 10 cm above the dye uptake. The dotted grid denotes the extensional region in b. The (b, c) East side at (b) 10 cm and (c) 30 cm above the dye uptake. Arrows show the border of the stained area. (d) Stained area near the heat sensor. Arrows show stained xylem. Scale bars: (b, c) 3 mm; (d) 4 mm.

The distribution of embolized vessels around the sensor hole was observed by means of cryo-SEM (Fig. 4). Embolisms were observed only within 2 cm above (Fig. 4a, b) and immediately below a hole. The number of water-filled vessels increased with increasing distance from the drill hole.

**Figure 4.**
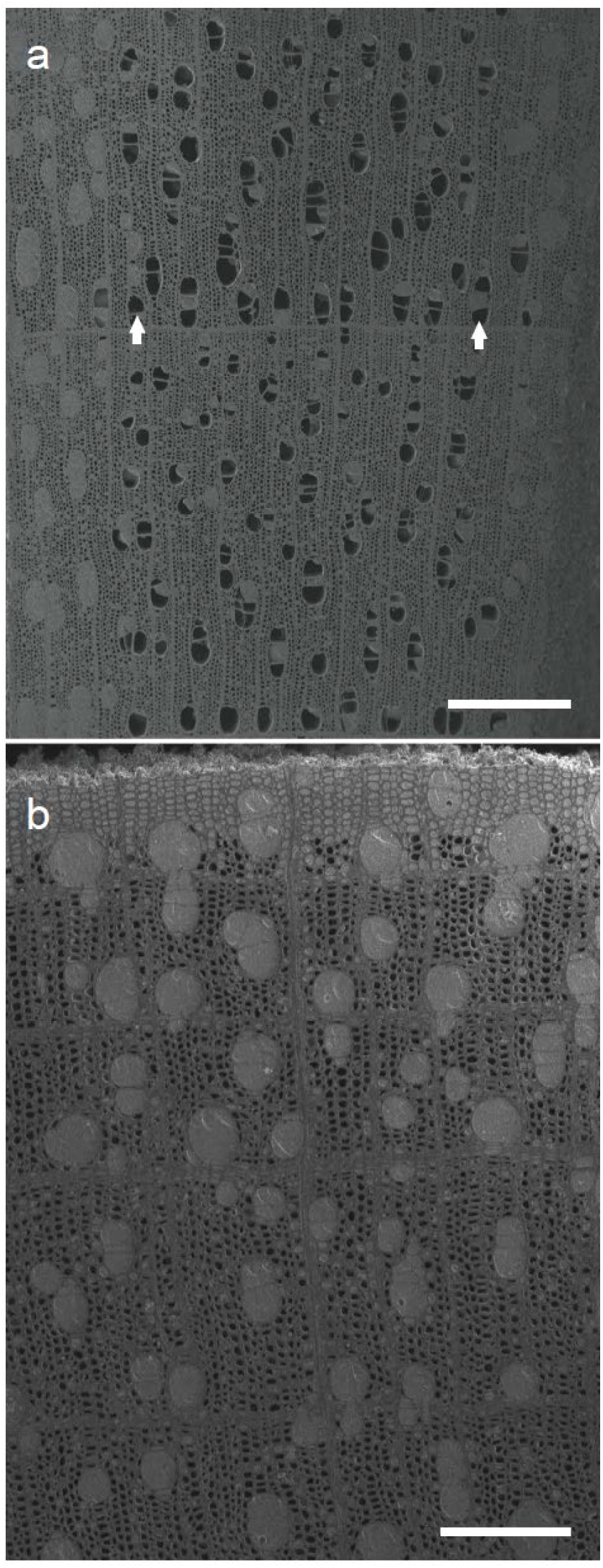
Cryo-SEM photos around holes drilled to insert sensors in *Betula ermanii*. (a) Just above and (b) 2 cm above a hole. Scale bars: (a) 500 μm; (b) 200 μm.

## Discussion

To understand the spatial distribution of water movement in intact stems, we monitored *F*_d_ before and after the creation of severed conduits below each TD sensor by drilling the stem under water. Just after drilling, we detected steep peaks in *F*_d_ in both trees. This steep peak is probably induced by a dramatic increase in water supply via severed conduits (Čermák et al. 1992). *F*_d_ via severed conduits and dye ascent showed similar velocities on each side (Table 1, Fig. 1c, d). In the results of Tsuruta et al. (2010) in *Chamaecyparis obtusa*, the dye heights were also 0.4 or 0.5 m h^–1^ in any sides (Fig. S2), although the circumferential variation of *F*_d_ values in intact trees was remarkably detected. Thus, it supported our results, suggesting that the circumferential variation may be induced in intact trees (Lu et al. 2000; Tateishi et al. 2008; Tsuruta et al. 2010), but the variation may be lost with limitless supply of water with the creation of severed conduits by drilling. This significant change of *F*_d_ values before and after drilling is potential source of error in an attempt to explain water movement in the stems of the mature trees.

On the other hand, Čermák et al. (1992) reported that the circumferential variation in the stem of Norway spruce was detected by dye uptake after one whole trunk was cut. The discordance between our results and the result of one tree in Čermák et al.’s (1992) report is still unknown, but the difference can arise between water uptake at the cut surface of the whole stem by saw and at a portion by drilling, or by growth conditions in the field such as partial shading of the tree crown (Oren et al. 1999). More detailed studies using both dye uptake and heat-transfer methods in the same tree are needed to understand intact water movement in mature trees.

The helical movement was clear with increasing distance from the dye uptake point (Fig. 3a). Although water movement by local diffusion which leads to a misrepresentation of the functional water-conducting pathway (Umebayashi et al. 2007) may be detected over 2 h, the stained time of this study (30 min) was much shorter. If helical movement widely occurs in intact trees, data collected by the heated transfer method may vary among sensor positions.

Accuracy tests of heat-transfer methods are conducted to confirm the technical reliability of various methods of measuring sap flow (e.g., Smith and Allen 1996; Clearwater et al. 1999; Steppe et al. 2010). In small-diameter samples, the methodology of the calibration is well discussed (Clearwater et al. 1999; Bush et al. 2010), and the estimation of the sap flow using TD method is also useful (Bush et al. 2010). On the other hand, in large stem, the TD method showed the largest underestimation among the three heat-transfer methods (Steppe et al. 2010). Steppe et al. (2010) suggested that TD method was not based on physical principles of heat transfer. In the TD sensor position, stained vessels were wholly distributed along the helical ascent (Fig. 3d), and the distribution of embolized vessels were also limited around the hole (Fig. 4). These results support that there is circumferential variation of water movement in the intact stem. Our results also suggest that the helical movement in the stem may be one of main factors on physical principles of water ascent, and the understanding leads to enhance the accuracy of *F*_d_ values using the TD method.

In summary, water movement via severed conduits after drilling treatments is different from water ascent in intact trees because of the loss of circumferential variation of sap flow in the stem. A simple comparison between *F*_d_ and the velocity of dye solution means little.

Future researches will need to involve collection of more spatial information about the role of *F*_d_ in the tangential water movement. Our results suggested that the development of non-invasive method and future technological advancement combining heated transfer and dye uptake methods need to understand the intact water movement in mature trees.

## Acknowledgments

This research was supported by the Japanese Ministry of Education, Culture, Sports, Science and Technology (MEXT) through a Grant-in-Aid for Scientific Research (#20780119). We thank Dr. Kenji Fukuda, The University of Tokyo, Dr. Hiroyuki Muraoka, Gifu University, and Dr. Kyoichi Otsuki, Kyushu University, for their technical support. We thank Mr. K. Kurumado (River Basin Research Center, Gifu University) for help felling the sample trees.

