## Supplemental Figure for "Spatial variations in sap flow rates in mature tree stems before and after drilling treatment"

**Supplementary material**

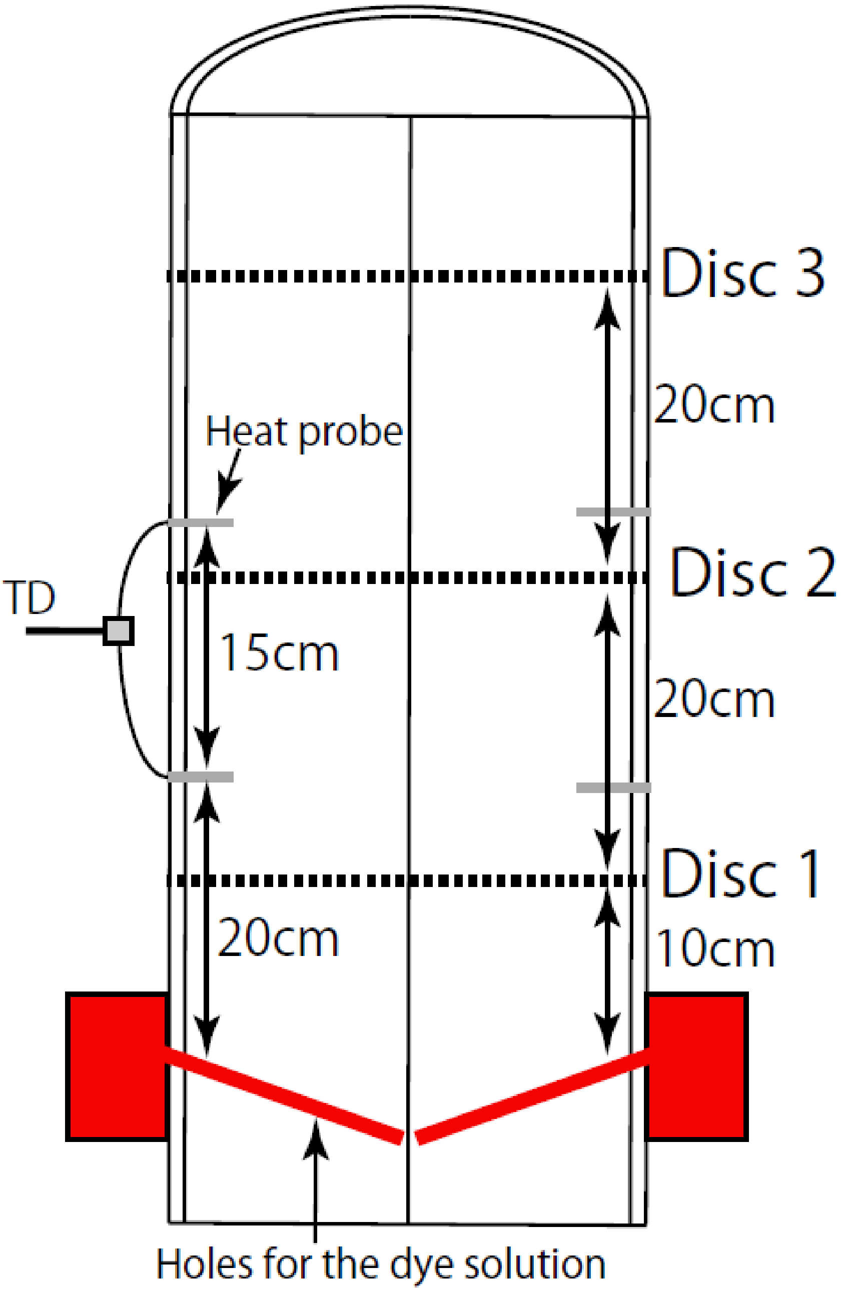


Figure S1. Schematic diagram of the thermal dissipation (TD) sensors and the dye uptake method in this study. Dash lines show the position to take the stained discs.

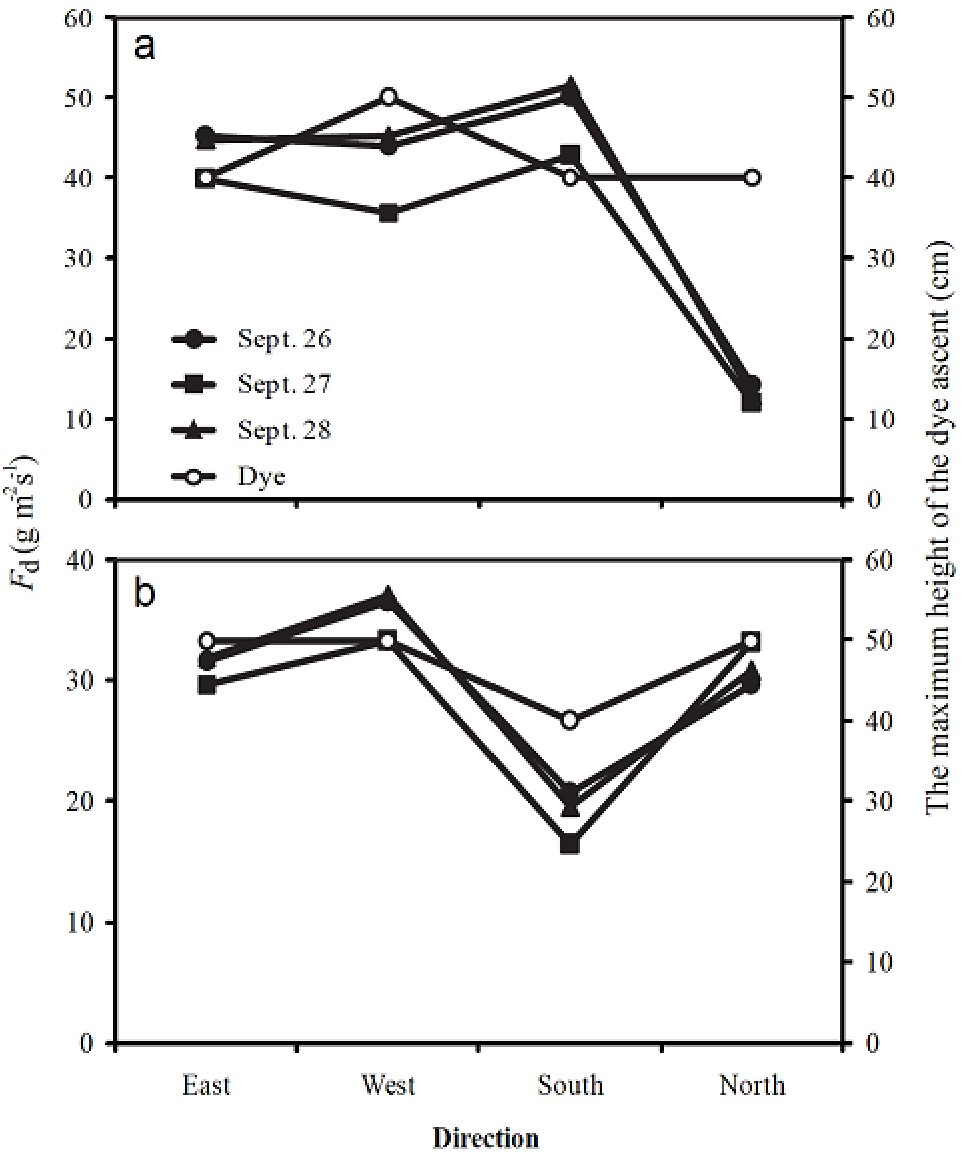


Figure S2. Midday patterns of mean sap flux density ($F_d$) and the maximum dye heights on each side of *Chamaecyparis obtusa* trees from 26 to 28 September, of trees (a) c1 and (b) c2. $F_d$ was calculated as the mean from 10:00 to 14:00 h. The dataset of both trees (c1 and c2) is the same as L1 and L3 in Tsuruta et al. (2010).
